# Structural basis of GTPase-mediated mitochondrial ribosome biogenesis and recycling

**DOI:** 10.1101/2021.03.17.435767

**Authors:** Hauke S. Hillen, Elena Lavdovskaia, Franziska Nadler, Elisa Hanitsch, Andreas Linden, Katherine E. Bohnsack, Henning Urlaub, Ricarda Richter-Dennerlein

## Abstract

Ribosome biogenesis is an essential process that requires auxiliary factors to promote folding and assembly of ribosomal proteins and RNA. In particular, maturation of the peptidyl transferase center (PTC), the catalytic core of the ribosome, is mediated by universally conserved GTPases, but the molecular basis is poorly understood. Here, we define the mechanism of GTPase-driven maturation of the human mitochondrial ribosomal large subunit (mtLSU) using a combination of endogenous complex purification, *in vitro* reconstitution and cryo-electron microscopy (cryo-EM). Structures of transient native mtLSU assembly intermediates that accumulate in GTPBP6-deficient cells reveal how the biogenesis factors GTPBP5, MTERF4 and NSUN4 facilitate PTC folding. Subsequent addition of recombinant GTPBP6 reconstitutes late mtLSU biogenesis *in vitro* and shows that GTPBP6 triggers a molecular switch by releasing MTERF4-NSUN4 and GTPBP5 accompanied by the progression to a near-mature PTC state. In addition, cryo-EM analysis of GTPBP6-treated mature mitochondrial ribosomes reveals the structural basis for the dual-role of GTPBP6 in ribosome biogenesis and recycling. Together, these results define the molecular basis of dynamic GTPase-mediated PTC maturation during mitochondrial ribosome biogenesis and provide a framework for understanding step-wise progression of PTC folding as a critical quality control checkpoint in all translation systems.

---

The human mitochondrial ribosome (mitoribosome) synthesizes thirteen essential subunits of the oxidative phosphorylation (OXPHOS) system. Defects in mitoribosome biogenesis or function result in OXPHOS deficiency and cause severe early-onset mitochondrial diseases^1^. The mitoribosome is composed of a small and a large ribosomal subunit (mtSSU and mtLSU, respectively), which each contain a ribosomal RNA (12S, 16S rRNA) and a number of mitoribosomal proteins (MRPs). Structural analyses of the mitoribosome have revealed its overall architecture and differences to cytoplasmic and bacterial ribosomes^2,3^. However, little is known about the mechanisms of mitoribosome assembly.

Biogenesis of the large ribosomal subunit (LSU) in all translation systems proceeds through distinct steps in which the formation of the peptidyl transferase center (PTC), the active site of the ribosome, represents the last and most critical step and requires the assistance of assembly factors^4–7^. In particular, universally conserved GTPases, which act as quality control and anti-association factors play key roles during late LSU assembly stages and ensure proper maturation of the PTC^8^. In human mitochondria, the GTPases GTPBP5, GTPBP6, GTPBP7 and GTPBP10 mediate mtLSU maturation, as their deletion results in a loss of active mitoribosomes and accumulation of incompletely assembled mtLSU intermediates^8–14^. These conserved GTPases act in concert with mitochondria-specific factors such as MALSU1, MTERF4 and NSUN4, to facilitate late steps of mtLSU maturation, but the mechanistic basis of GTPase-driven LSU biogenesis and PTC folding is not known.

## Two distinct mtLSU biogenesis intermediates accumulate in the absence of GTPBP6

To investigate this, we isolated mtLSU complexes from a human cell line lacking GTPBP6, which has been shown to accumulate assembly intermediates that contain GTPBP7, GTPBP10, and GTPBP5 as well as MALSU1, MTERF4 and NSUN4^12^. Mass-spectrometric analyses confirmed the presence of these proteins as well as all 52 MRPs (**Extended Data Fig. 1a, Supplementary Table 1**). We then analyzed the intermediates by single-particle cryo-EM (dataset 1). Particle classification yielded two distinct reconstructions of mtLSU assembly intermediates at overall resolutions of 2.2 Å and 2.5 Å, respectively, which led to refined structures (**Extended Data Fig. 1b-e, Extended Data Table 1**).

Compared to the mature mtLSU, both reconstructions show an extra density close to the L1 stalk and a distinct folding of the interfacial rRNA (**Fig. 1a**). The density could be unambiguously fit with the crystal structure of the RNA-binding protein MTERF4 and the methyltransferase NSUN4^15,16^, with some adjustments (**Fig. 1a, Extended Data Fig. 2a**). Additionally, both structures contain all MRPs as well as the MALSU1-L0R8F8-mtACP module, which prevents premature subunit association ^5^. However, they differ in their PTC conformation, the second reconstruction showed an additional density above the PTC. Based on its resemblance to bacterial Obg^17^, we identified it as GTPBP5 (**Fig. 1a, Extended Data Fig. 2b**). Thus, two distinct mtLSU biogenesis intermediates accumulate in the absence of GTPBP6, one containing MTERF4-NSUN4 and one additionally containing GTPBP5.

**Fig. 1.**
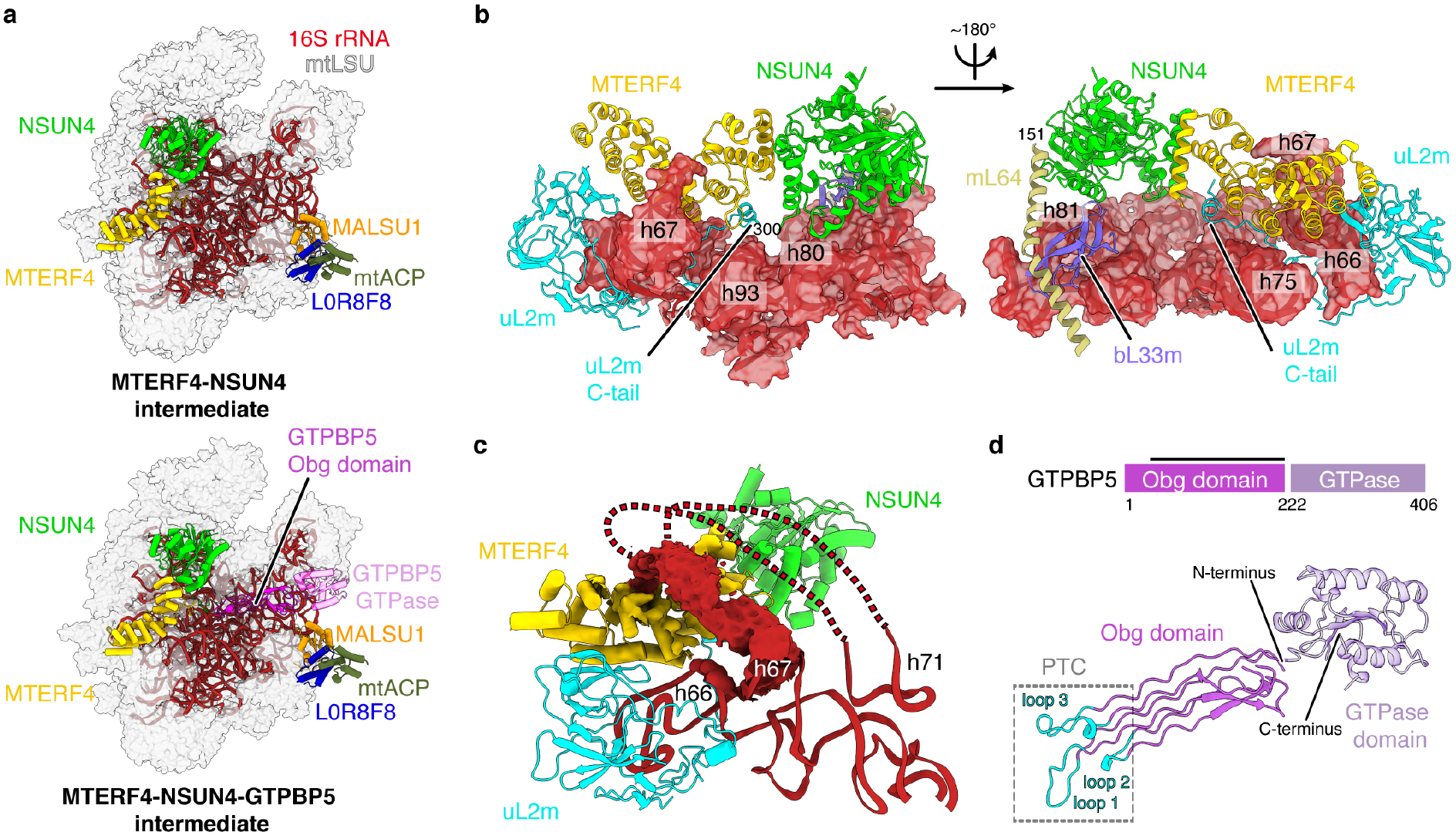
Structure of the MTERF4-NSUN4 and MTERF4-NSUN4-GTPBP5 bound mtLSU assembly intermediates. (a) Cryo-EM structure of mtLSU assembly intermediates. The 16S rRNA (red) and indicated biogenesis factors are shown as cartoon and the remaining MRPs are shown as grey transparent surface. (b) Interaction of the MTERF4-NSUN4 complex with the mtLSU. Regions of the 16S rRNA interacting with MTERF4-NSUN4 are shown as cartoon and as transparent surface. (c) The region encompassing h68 – h71 of the 16S rRNA wraps above MTERF4 as indicated by red lines. The density is shown as red surface. (d) Structure of human GTPBP5. The protein is depicted schematically on top, with domain annotation indicated by coloring and residue numbers. The black bar above represents regions modeled in the structure. A homology model of the GTPase domain is shown transparently. The conserved loops 1-3 that contact bases of the PTC are shown in cyan.

## MTERF4 binds subunit-bridging elements in the rRNA

The structure of the MTERF4-NSUN4-bound intermediate shows that this complex binds to the interfacial side of the 16S rRNA (**Fig. 1a**). Compared to previous crystal structures, the curvature of MTERF4 is altered by an inward rotation of helices α1-8, resulting in a narrower RNA binding groove (**Extended Data Fig. 2a**). MTERF4 interacts with the mtLSU through contacts with h75 in the L1 stalk (nucleotides 2743-2756 and 2792-2804) and with the C-terminal tail of uL2m (residues 275-300) (**Fig. 1b**). In the mature mtLSU, this tail runs in between h68 and h66^3^. In the MTERF4-bound state, it is rearranged beneath MTERF4 and forms a helix (residues 290-295) that binds α12 of MTERF4. NSUN4 binds to the mtLSU on top of the P loop/h80 (nucleotides 2815-2821) and h81 (nucleotides 2841-2853) (**Fig. 1b**), next to bL33m. The N-terminal part of mL64 runs above NSUN4, but this region is invisible, as in previous structures^3^. The active site of NSUN4 is positioned above h81, but the closest RNA base is more than 15 Å away from its SAM-binding site^15,16^, suggesting that NSUN4 does not methylate the 16S rRNA.

MTERF4 additionally contacts the interfacial rRNA segment that forms h68-h70 in the mature mtLSU. In the MTERF4-NSUN4 assembly intermediate, this region is partially unfolded and wraps over the outward-facing RNA-binding groove of MTERF4 (**Fig. 1c**). Although the density did not allow for atomic modeling, comparison shows that helices h68 and h69 would clash with MTERF4 in their mature conformation (**Extended Data Fig. 2c**), indicating that MTERF4 binds a premature conformation of the interfacial rRNA. In the mature mitoribosome, h68-h70 form seven of the fifteen intersubunit bridges^2^. MTERF4-NSUN4 may thus act as a quality control checkpoint by sequestering the interfacial rRNA to prevent subunit joining prior to final mtLSU maturation.

## GTPBP5 and NSUN4 cooperate to facilitate PTC folding

The structure of the second mtLSU intermediate reveals the structure and function of GTPBP5 (**Fig. 1a**). GTPBP5 binds above the PTC and interacts primarily with the rRNA (**Extended Data Fig. 2d**). It consists of an N-terminal Obg domain and a C-terminal GTPase domain (**Fig. 1d**). The GTPase domain is poorly resolved, but the density indicates that it resides between uL11m and the sarcin-ricin loop (SRL, h95). The Obg domain extends along the PTC and contains three conserved loops at its tips (loop 1: residues 93-103, loop 2: residues 140-145, loop 3: residues 191-205)^17^ (**Fig. 1d)**. These loops reach into the PTC, where we additionally observe density for the N-terminal tail of NSUN4 (residues 26-37), which was invisible in the intermediate lacking GTPBP5 but becomes ordered in the presence of GTPBP5 (**Fig. 2**).

**Fig. 2.**
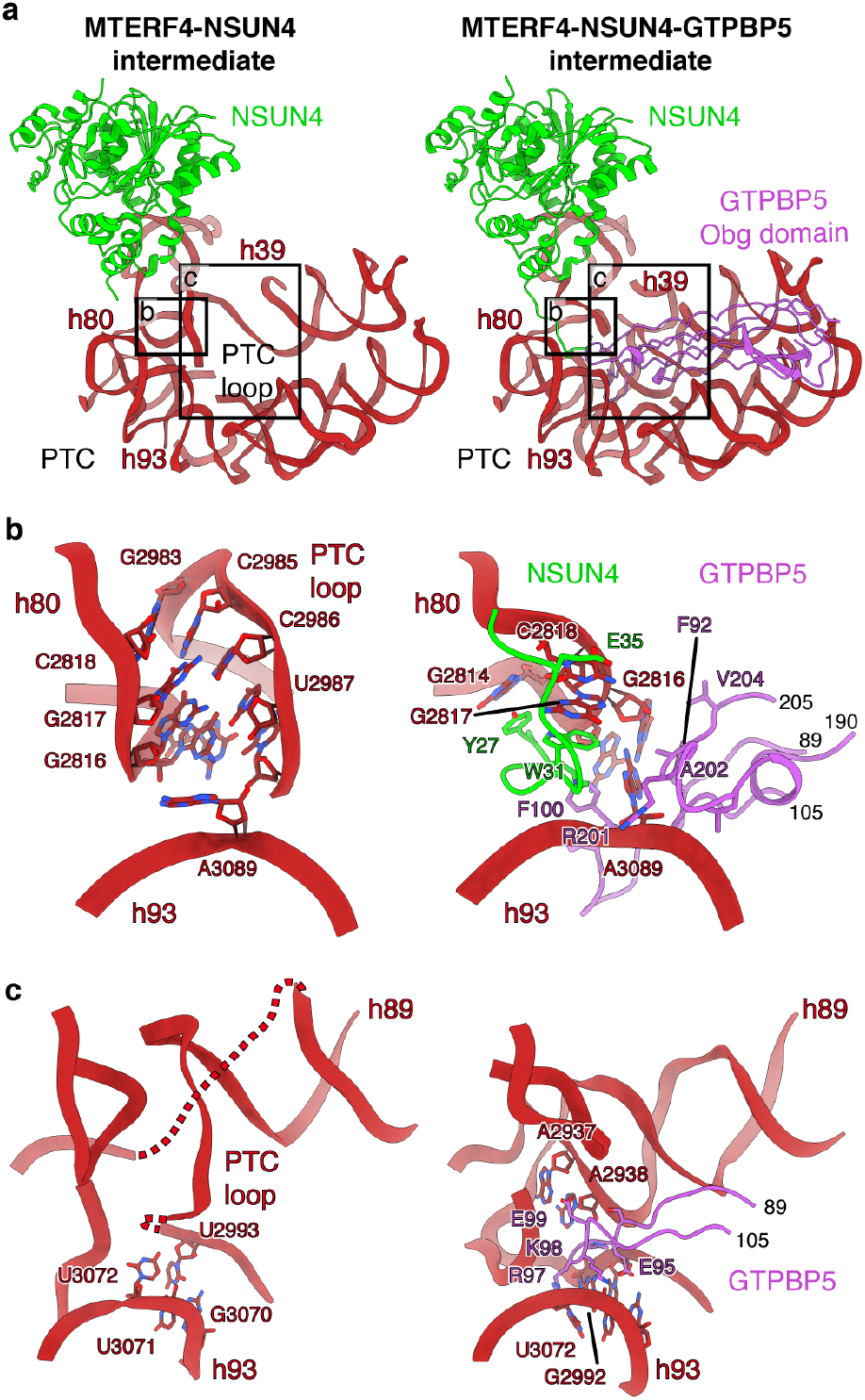
GTPBP5 cooperates with NSUN4 to mature the PTC. **(a)** Close-up view of the PTC in the NSUN4-MTERF4 (left) and NSUN4-MTERF4-GTPBP5 (right) mtLSU assembly intermediates. **(b, c)** Re-arrangements in the P loop/h80, h93 and PTC loop upon GTPBP5 binding. Selected bases that undergo rearrangements are indicated and highlighted as sticks. Loop 1 and 3 of GTPBP5 and the N-terminal tail of NSUN4 are shown as cartoons and residues that interact with the indicated bases are shown as sticks.

Comparison of the two assembly intermediates shows that they differ in their PTC maturation state. In the intermediate lacking GTPBP5, the PTC is partially disordered and adopts a premature conformation (**Fig. 2a**). In particular, h72 (2692-2695), the PTC loop (nucleotides 2936-2946 and 2977-2992), h39 (nucleotides 2108-2115) and the P loop/h80 (nucleotides 2814-2818) adopt distinct conformations. The PTC loop is partially mobile, and nucleotides 2975-2995 form a loop that base pairs with the tip of the P loop/h80 (**Fig. 2b, c**). In the GTPBP5-containing intermediate, the PTC adopts a more mature-like conformation in which the PTC loop forms the base of h89, and the h80 tip is shifted upward (**Fig. 2b**). The structures show how GTPBP5 and the NSUN4 tail facilitate these rearrangements. First, GTPBP5 and the NSUN4 tail disrupt the interaction between the PTC loop and h80, which allows the PTC loop to refold. NSUN4 sequesters bases G2817 and G2814 through stacking interactions with W31 and Y27, respectively (**Fig. 2b**), and GTPBP5 binds G2816 through stacking interactions with F92 and sandwiches A3089 between F100, A202 and F92. Second, GTPBP5 stabilizes the refolded PTC conformation through backbone and base interactions with charged residues in loop 1 (R97, K98, E95 and E99) (**Fig. 2c**). These rearrangements also lead to ordering of ribosomal protein elements which are disordered in the intermediate without GTPBP5, including parts of mL63 (residues 9-21), uL10m (residues 30-36) and uL16m (residues 47-69 and 134-148).

PTC maturation also involves 2’-*O*-methylation of bases within the P loop/h80 (G2815) and the A loop/h92 (U3039 and G3040) catalyzed by MRM1, MRM2 and MRM3, respectively^14,18,19^. Methylation of U3039 is one of the last steps in PTC maturation and requires GTPBP5^14^. The cryo-EM reconstructions show density consistent with 2’-*O*-methylation at all these residues (**Extended Data Fig. 2e**), suggesting that GTPBP6 acts downstream of MRM1-3 and GTPBP5, in agreement with previous data^12^.

Taken together, the cryo-EM structures reveal how GTPBP5 and NSUN4 facilitate maturation of the PTC and suggest a sequential sequence of PTC methylation and folding (**Supplementary Video 1**). They also explain the dual-role of NSUN4 as a methyltransferase in mtSSU assembly^20^ and as a biogenesis factor in mtLSU maturation, as its N-terminal tail is not conserved in the homologous bacterial methyltransferase rsmB.

## GTPBP6 displaces GTPBP5 and MTERF4-NSUN4

We next aimed to investigate the role of GTPBP6 during mtLSU biogenesis. For this, we reconstituted mtLSU maturation *in vitro* by complementing the biogenesis intermediates from GTPBP6-deficient cells with recombinant GTPBP6 in the presence of GTP and ATP. Subsequent cryo-EM analysis again revealed the MTERF4-NSUN4- and the MTERF4-NSUN4-GTPBP5-bound mtLSU complexes (dataset 2; **Extended Data Fig. 3, Extended Data Table 2**), which were largely identical as before but showed improved density for the interfacial rRNA and the GTPBP5 GTPase domain (see experimental procedures) (**Extended Data Fig. 4a-d**). In addition, classification yielded a 2.6 Å reconstruction that lacks MTERF4-NSUN4 and GTPBP5 but shows a new density that corresponds to GTPBP6 in close proximity to the L12 stalk, which allowed us to build a molecular model (**Fig. 3a,b, Extended Data Fig. 3, 4e**). GTPBP6 contains a N-terminal nucleotide-binding domain (NTD), a PTC-binding linker domain, a GTPase domain and a C-terminal domain (CTD) (**Fig. 3b**). Like GTPBP5, it binds above the PTC and primarily interacts with rRNA (**Extended Data Fig. 4f**). The NTD stacks against h71, which appears stabilized compared to the MTERF4-NSUN4 state. The PTC-binding domain resides between h89 and h92, and inserts a loop (residues 241-255) deep into the PTC. The GTPase domain is located next to the NTD, on top of h89, and shows clear density for a bound GTP molecule, suggesting that GTP hydrolysis is not required for GTPBP6 binding (**Fig. 3b, Extended Data Fig. 4g**). The CTD is positioned between the SRL (h95) and uL11m, in a similar position as the GTPase domain of GTPBP5 (**Extended Data Fig. 4f,h**).

**Fig. 3.**
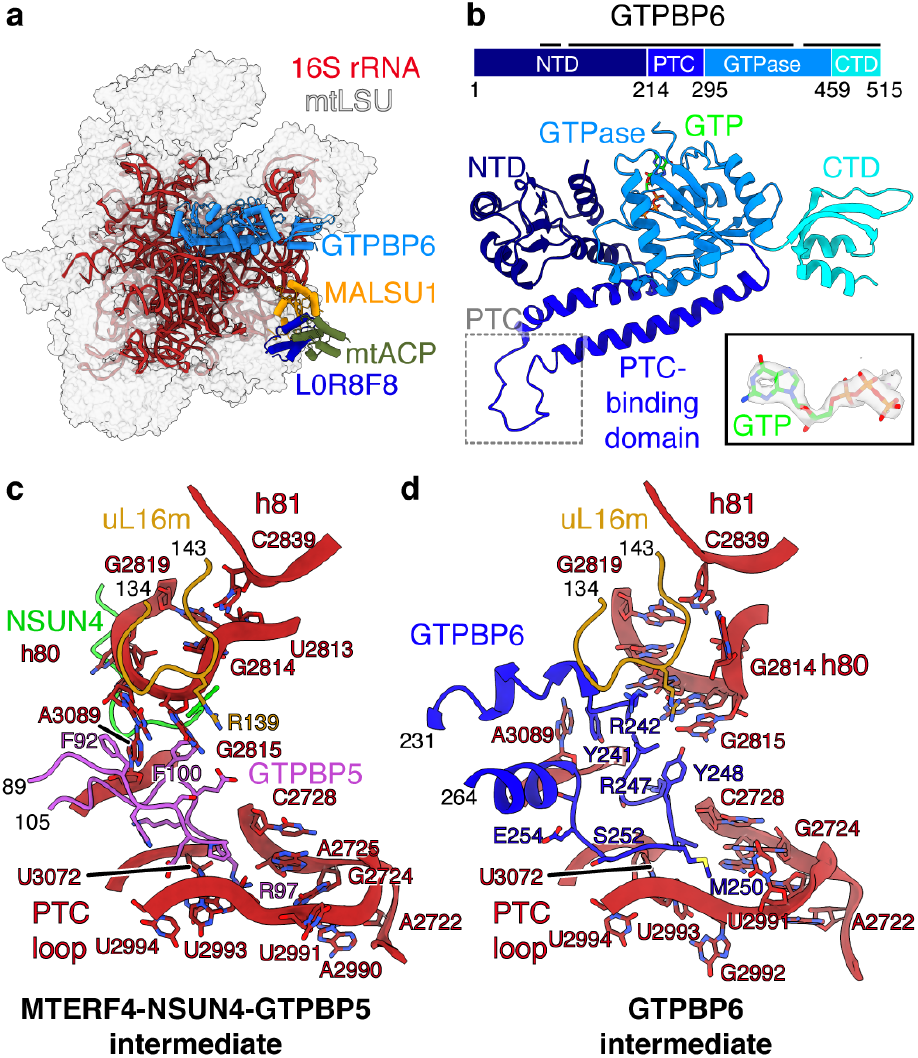
GTPBP6 binding causes rearrangements in the PTC. **(a)** Cryo-EM structure of the GTPBP6-bound mtLSU assembly intermediate. **(b)** Structure of human GTPBP6. The protein is depicted schematically on top, with domain annotation indicated by coloring and residue numbers. The black bar above represents regions modeled in the structure. The density for the bound GTP is shown as indent. **(c, d)** Re-arrangements in the PTC upon GTPBP6 binding. The PTC region is shown enlarged in the MTERF4-NSUN4-GTPBP5 (c) and the GTPBP6 (d) mtLSU assembly intermediates.

Comparison of the GTPBP6- and GTPBP5-bound structures suggests a mechanism for the hierarchical action of these factors during mtLSU biogenesis. Superimposition shows that both factors share the same binding site, and that the NTD of GTPBP6 would clash with NSUN4 (**Extended Data Fig. 4h**). Thus, binding of GTPBP6 and GTPBP5 / MTERF4-NSUN4 is mutually exclusive and GTPBP6 triggers the release of these factors from the mtLSU.

## GTPBP6 mediates PTC maturation

Comparison to the GTPBP5-bound state also reveals that GTPBP6 further folds the PTC. The PTC-binding loop of GTPBP6 occupies the same position as loop 1 of GTPBP5, between the P loop/h80 and the PTC loop and in vicinity to uL16m (residues 134-143), and causes rearrangements in the elements (**Fig. 3c,d, Supplementary Video 1**). In particular, the tip of the P loop/h80 (residues 2814-2819) refolds, which leads to elimination of a base pair between G2819 and U2813 and replacement of the latter by C2839 from h81. G2814 is flipped out of the loop to face outward, and G2815 is rearranged and may contact Y248 in GTPBP6 and R139 in uL16m.

Binding of GTPBP6 also leads to movement of A3089 by replacing its interaction with F92 in GTPBP5 by a stacking interaction with Y241 in GTPBP6. Finally, nucleotides 2990-2994, 2722-2728 and U3072 in the PTC loop undergo conformational changes, which may be induced by a change in chemical environment upon exchange of maturation factors, as GTPBP6 places hydrophobic residues (L249, M250) at the position previously occupied by R97 of GTPBP5. Collectively, these rearrangements lead to a nearly-mature PTC conformation.

## A molecular model of GTPase-mediated PTC maturation

These structural snapshots allow us to deduce a model for GTPase-mediated mtLSU assembly and PTC maturation, which we have summarized in a molecular movie (**Extended Data Fig. 5, Supplementary Video 1**). First, early biogenesis factors assemble a core mtLSU with a premature PTC and partially unfolded interfacial rRNA that lacks late-stage MRPs as well as 2’-*O*-methylations within the PTC^14^. This mtLSU intermediate contains the MALSU1-L0R8F8-mtACP module and the MTERF4-NSUN4 complex, which both prevent premature subunit joining^5^. PTC methylations are then introduced by MRM1 and MRM2, leading to the intermediate with methylated but unfolded PTC. Next, GTPBP5 and NSUN4 act in concert to establish the basic architecture of the PTC. Both GTPBP5 and the MTERF4-NSUN4 are then replaced by GTPBP6, which induces further conformational changes that lead to a near-mature PTC. The release of MTERF4-NSUN4 liberates the rRNA region h68-h71, which can then adopt its final conformation to facilitate subunit joining. Final maturation steps must then involve dissociation of GTPBP6 and the MALSU1-L0R8F8-mtACP module to allow formation of the functional 55S mitoribosome.

## Mechanism of GTPBP6-mediated ribosome recycling

In addition to its role in mtLSU biogenesis, GTPBP6 also facilitates the dissociation of intact 55S mitoribosomes into subunits, which might be required to rescue stalled ribosomes^12^. To determine the mechanism of GTPBP6-mediated ribosome splitting, we treated 55S mitoribosomes with recombinant GTPBP6 in the presence of GTP and ATP. Subsequent cryo-EM analysis revealed the presence of complete mitoribosomes as well as free mtLSU and mtSSU particles (**Extended Data Fig. 6, Extended Data Table 3**). Classification of the mtLSU particle population led to a reconstruction at an overall resolution of 2.7 Å, which shows that GTPBP6 binds to the mature mtLSU in the same location as during ribosome biogenesis (**Fig. 4a,b**). In addition to the previously observed contacts, GTPBP6 also interacts with h69 in the mature mtLSU (**Fig. 4c**), which was unfolded in the biogenesis intermediates but forms intersubunit contacts with the mtSSU in the 55S mitoribosome. The NTD of GTPBP6 binds to h69 and inserts a tryptophan residue (W107) next to the U2575-A2582 basepair at its tip. Superimposition with the intact mitoribosome shows that h69 is shifted by ~7 Å in the GTPBP6-bound state, which leads to clashes with h44 in the mtSSU (**Fig. 4d**). Thus, GTPBP6 dissociates the ribosome by rearranging elements that mediate intersubunit interactions, as has been suggested for HflX and ribosome recycling factors^21–24^. However, its mechanism is distinct, because W107 is not conserved and HflX does not form direct contacts with bases in h69, yet causes a more prominent displacement (13 Å vs 7 Å)^23^.

**Fig. 4.**
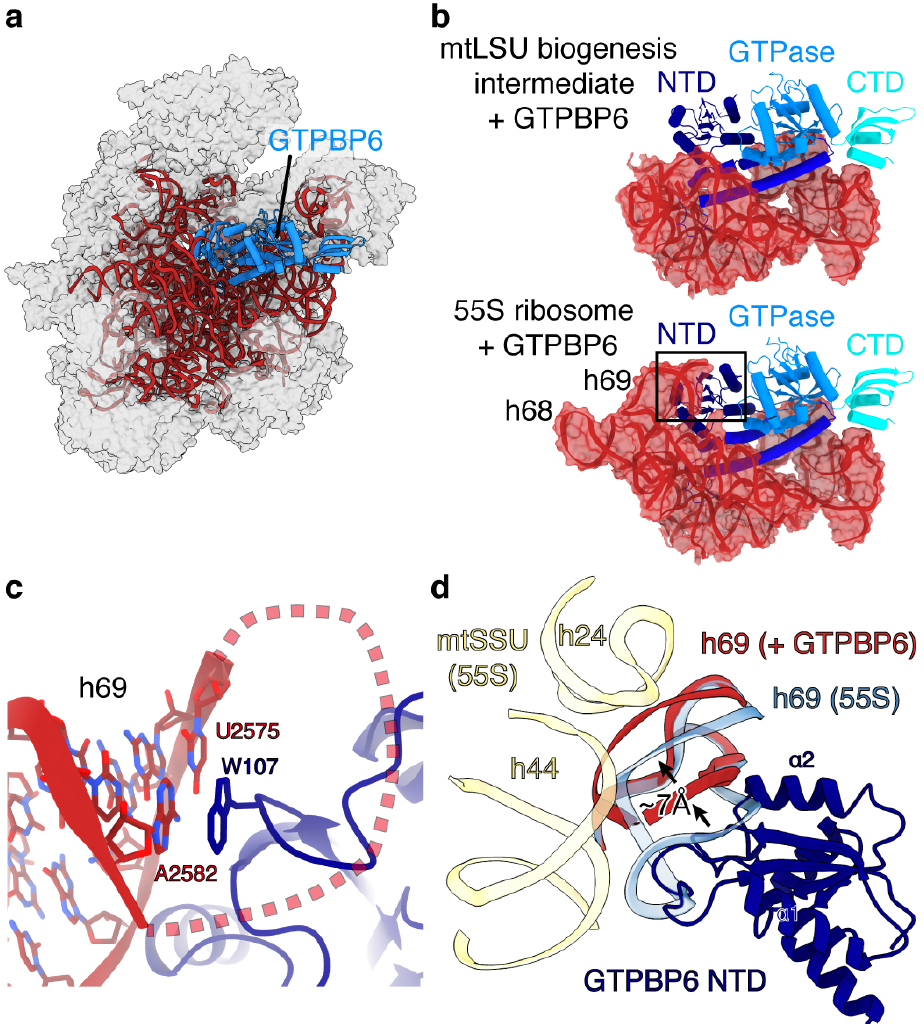
Structural basis of ribosome recycling by GTPBP6. **(a)** Cryo-EM structure of GTPBP6 bound to the mature mtLSU. **(b)** Comparison of 16 rRNA interactions of GTPBP6 during biogenesis (top) and ribosome splitting (bottom). **(c)** GTPBP6 interacts with h69. Close-up view of h69 in the split mature mtLSU bound to GTPBP6. W105, which forms stacking interactions with h69, is shown as sticks. **(d)** Comparison of intersubunit-bridging elements in the 55S mitoribosome and the GTPBP6-bound split mtLSU. The GTPBP6-bound split mtLSU structure was superimposed with the structure of the elongating 55S ribosome (PDB 6ZSG)^31^. h69 is shown in blue (55S mitoribosome) or red (GTPBP6-bound mtLSU). GTPBP6-binding causes a shift of h69 by approximately 7 Å leading to clashes with h24 and h44 (yellow) in the mtSSU.

The structure of GTPBP6 bound to the mature mtLSU also reveals two distinct conformations of the PTC binding loop (**Extended Data Fig. 7a**). While the first is identical to that observed during mtLSU biogenesis, the second shows rearrangements in α7 and α8 and a register in α5 that translates into the PTC. This is accompanied by rearrangements of PTC bases, including A3089, U3072 and U2993, which play key roles during peptide bond formation and peptide release ^25–29^ and undergo conformational changes during elongation and termination in bacteria ^27^,^28^ (**Extended Data Fig. 7b**). However, it is not clear whether GTPBP6 induces these rearrangements or recognizes different PTC configurations that occur during the peptide elongation cycle.

The preferred substrates of GTPBP6 are vacant ribosomes or post-hydrolysis complexes with a deacylated tRNA in the P site^12^. Our structural data explain this preference, as a peptidyl tRNA in the P site would prevent GTPBP6 binding. To recycle ribosomes, release factors that trigger peptide hydrolysis such as ICT1/mL62 must therefore act prior to GTPBP6, as has been suggested recently^12^,^23^,^30^. Alternatively, GTPBP6-mediated ribosome recycling may require spontaneous hybrid P/E state formation, in agreement with *in vitro* data^12^.

In summary, these data provide the structural and mechanistic basis of mtLSU maturation by showing how universally conserved and mitochondria-specific assembly factors act in concert to mediate the step-wise folding of the PTC. This provides the framework for understanding ribosome biogenesis in general, as GTPase-driven PTC maturation is conserved throughout all domains of life.

## ACKNOWLEDGEMENTS

This work was funded by the Deutsche Forschungsgemeinschaft by the Emmy-Noether grant [RI 2715/1-1 to R.R.-D.]; the Excellence Cluster [EXC 2067/1-390729940 to R.R.-D. and H.S.H]; the Forschergruppe 2848 [to H.S.H.], the Collaborative Research Center [SFB860 to R.R.-D., K.E.B and H.U.; SFB1160 to H.S.H.]; and the Max Planck Society [to H.U.].

## AUTHOR CONTRIBUTIONS

H.S.H. prepared samples for cryo-EM, collected and processed cryo-EM data, built and interpreted structural models. E.L. and F.N. isolated ribosomal complexes from human cells. E.H. established ribosome purification strategy. A.L. and H.U. performed mass spectrometry analysis. K.E.B. performed the bisulfite sequencing. H.S.H., E.L. and R.R.-D. prepared figures. H.S.H. and R.R.-D. wrote the manuscript.

## DATA AVAILABILITY

The electron density reconstructions and structure coordinates were deposited with the Electron Microscopy Database (EMDB) under accession code EMD-xxxx and with the Protein Data Bank (PDB) under accession code xxxx.

## EXPERIMENTAL PROCEDURES

No statistical methods were used to predetermine sample size. The experiments were not randomized, and the investigators were not blinded to allocation during experiments and outcome assessment.

### Cell culture conditions

HEK293-Flp-In T-Rex wild type (WT) (Thermo Fisher Scientific) and *Gtpbp6*^−/−^ cell lines were grown under standard cultivation conditions ^12^. Briefly, cells were cultured in Dulbecco’s modified Eagle’s medium (DMEM) with supplements (10% FBS (Sigma), 2 mM L-glutamine (GIBCO), 1 mM sodium pyruvate (GIBCO), 50 μg/ml uridine (Sigma-Aldrich)) in 5% CO_2_ humidified atmosphere at 37°C.

### Mitoplasts isolation

Mitoplasts isolation was performed essentially as described recently ^11^ with minor changes. In brief, cells were harvested and homogenized in trehalose buffer (300 mM trehalose, 10 mM KCl, 10 mM HEPES-KOH pH 7.4) with addition of 1 mM PMSF and 0.2% BSA using Homogenplus Homogenizer (Schuett-Biotec, Germany). After each homogenization step mitochondria were separated from cell debris and nuclei at 1000 xg for 10 min, 4°C. Obtained mitochondria were pelleted for large-scale mitoplast preparation. To isolate mitoplasts, mitochondria were subjected to digitonin/proteinase K treatment (detergent to protein ratio 1:4, proteinase K to protein ratio 1:200), washed 4 times with trehalose buffer and pelleted at 25 000 xg for 15 min, 4°C (SS34 Rotor, Beckman Coulter).

### Purification of mitoribosomes

Mitoplasts were lysed in lysis buffer (20 mM Tris-HCl (pH 7.4), 100 mM NH_4_Cl, 15 mM MgCl_2_, 2 mM DTT, 1% Triton X-100) with detergent to protein ratio 2.5:1 and the resulting lysate was clarified by centrifugation at 16 000 xg for 15 min, 4°C. To enrich mitoribosomal particles and to eliminate other mitochondrial protein complexes contaminants, the collected supernatant was subjected to a two-step sucrose cushion (1 M sucrose cushion/1.75 M sucrose cushion) and centrifuged for 15 h at 148 000 xg at 4°C. Fractions were collected from top to bottom of the cushion and the fraction containing mitoribosomal particles was concentrated and subsequently washed with 5 volumes of wash buffer (100 mM NH_4_Cl, 15 mM MgCl_2_, 20 mM Tris-HCl (pH 7.4), 2 mM DTT) in order to reduce the sucrose concentration and the sample volume. Concentrated sample was loaded on a 15-30% sucrose gradient (15-30% (w/v) sucrose, 100 mM NH_4_Cl, 15 mM MgCl_2_, 20 mM Tris-HCl (pH 7.4)), centrifuged at 115 600 xg for 16 h 10 min, 4°C (SW41Ti rotor, Beckman Coulter) and 16 fractions were collected. Each fraction was measured at 260 nm and fractions corresponding to the mtLSU or 55S mitoribosome were further concentrated and washed with wash buffer as described above. Purified mitoribosomal complexes were stored at - 80°C, or used directly for grid preparation or for *in vitro* reconstitutions followed by cryo-EM analyses.

To monitor the structural rearrangements in the mtLSU upon GTPBP6 binding 0.1-0.18 μM of 39S/ 55S particles purified from *Gtpbp6*^−/−^ or WT cell lines were mixed with 20-fold molar excess of purified GTPBP6 protein in the presence of nucleotides (1 mM ATP/1mM GTP) in reaction buffer (2 mM DTT, 100 mM NH_4_Cl, 15 mM MgCl_2_, 20 mM Tris-HCl (pH 7.4)). Mixtures were incubated at 4°C for 30 min and were used directly for grid preparation.

### Cryo-EM sample preparation, data collection and processing

Purified mtLSU or 55S mitoribosome samples (4 μL) were applied to freshly glow discharged R 3.5/1 holey carbon grids (Quantifoil) that were precoated with a 2-3 nm carbon layer using a Leica EM ACE600 coater. Prior to flash freezing in liquid ethane, the samples were incubated on the grid for 30 s in a Vitrobot MarkIV (Thermo Fisher Scientific) at 4°C and 100 % humidity and subsequently blotted for 3 s seconds with a blot force of 0. Cryo-EM data collection was performed with SerialEM ^32^ using a Titan Krios transmission electron microscope (Thermo Fisher Scientific) operated at 300 keV. Images were acquired in EFTEM mode with a slit width of 20 eV using a GIF quantum energy filter and a K3 direct electron detector (Gatan) at a nominal magnification of 81,000x corresponding to a calibrated pixel size of 1.05 Å/pixel. Exposures were recorded in counting mode with a dose rate of ~20 e^-^/px/s resulting in a total dose of 36 - 40 e^-^/Å^2^ (see **Extended Data Table 1-3**) that was fractionated into 40 movie frames. Motion correction, CTF-estimation, particle picking were performed on the fly using Warp ^33^. Particle extraction was performed with Relion ^34^ (dataset 1; 3-fold binned) or with Warp (dataset 2, dataset 3; unbinned).

For dataset 1 and dataset 2, particles were subjected to 2D classification followed by consensus 3D refinement using ab initio model created in cryoSPARC ^35^ as reference. Particles were then subjected to 3D classification without image alignment followed by unbinned reextraction (dataset 1) and 3D classification using a soft mask around the interfacial rRNA region where extra density for factors was visible (**Extended Data Fig. 1, 3**). Particle subsets were then subjected to CTF refinement (dataset 1 and 2) and Bayesian polishing (dataset 1). Further separation of particle populations with differing PTC conformations or subtle conformational differences within factors was achieved by using soft masks and a regularization parameter of T = 100 in Relion. Final maps were obtained by gold-standard 3D refinement followed by postprocessing in Relion. Focused refinements using soft masks were used to obtain improved maps for conformationally flexible regions. For dataset 3, 2D classification and initial further steps were carried out in cryoSPARC (**Extended Data Fig. 6**). Good classes were selected and a subset of 150,000 particles was used to generate three ab-initio models, which clearly resembled the full 55S mitoribosome, the mtLSU and the mtSSU. Particles were classified into these three classes by supervised classification in cryoSPARC and the mtLSU particle subset subject to consensus 3D refinement in Relion using the model from cryoSPARC as reference. Particles were further classified without image alignment, followed by focused classification with a soft mask around the interfacial rRNA and focused classification using a mask around GTPBP6 and T = 100. This led to three particle subsets containing GTPBP6. The first two represented different PTC conformations and were refined to high resolution, while the third resembled PTC conformation 1 but lacked clear density for h69. Final maps were obtained by gold-standard 3D refinement followed by postprocessing in Relion. Local resolution mas estimated using Relion. Figures were prepared with Chimera ^36^ and ChimeraX^37^.

Particle classification yielded two classes that resemble the mtLSU assembly intermediates previously observed in wild type cells, which both contain the MALSU1-L0R8F8-mtACP module and differ in their rRNA folding state and the presence of bL36m ^5,14^.

### Model building and refinement

An initial model for the MTERF4-NSUN4 intermediate was obtained by rigid body fitting the previously reported structure of a mtLSU assembly intermediate (PDB: 5OOL) ^5^ and the previously reported crystal structure of the MTERF4-NSUN4 complex (PDB 4FZV) ^16^ into the density in Chimera. The model was manually adjusted and rebuilt in Coot ^38^ and served as a starting model for the remaining structures. The model of GTPBP5 was generated by docking a homology model generated with SwissModel ^39^ into the density followed by manual rebuilding in Coot. The GTPase domain of GTPBP5 showed poor density and only allowed for docking and adjusting of the homology mode and was therefore omitted from the final model. The final model contains residues 74-161 and 175-222 of GTPBP5. The model of GTPBP6 was generated by docking a homology model generated by SwissModel into the density followed by manual rebuilding in Coot. The final model of GTPBP6 comprises residues 94-515, but residues 106-118, 424-431 and the loop contacting h69 (106-118) are invisible in the mtLSU biogenesis intermediate containing GTPBP6. The models were refined using phenix.real_space_refine ^40^ with restraints for 2’-methyl-UMP generated by phenix.elbow ^41^. Model quality was assessed with MolProbity ^42^. Figures were generated with ChimeraX.

Notably, the reconstructions obtained from dataset 1 lack strong density for uL24 and bL34m. We speculate this may be a consequence of grid preparation, as both proteins show strong density in the reconstructions from dataset2. In addition, we observed some differences between the MTERF4-NSUN4 and MTERF4-NSUN4-GTPBP5 intermediates from the first and the second dataset. First, the rRNA wrapping over MTERF4 is better resolved in dataset 2, showing that it forms a helical structure (**Extended Data Fig. 4a**). This is supported by basic residues in MTERF4 that form a charge-complementary binding groove, as well as two potential base-binding pockets (**Extended Data Fig. 4b**). Second, a part of the uL2m C-tail (residues 270-284) occupies a different path than in dataset1 (**Extended Data Fig. 4c**). Third, the GTPase domain of GTPBP5 is better resolved in dataset 2, which allowed rigid-body fitting of a homology model (**Extended Data Fig. 4d**).

### Expression and purification of human GTPBP6

Human Δ43GTPBP6 was cloned into the pET-derived vector 14-C (gift from Scott Gradia; Addgene plasmid #48309; http://n2t.net/addgene:48309; RRID:Addgene_48309) via ligation-independent cloning (primer sequences: for: (5’-TACTTCCAATCCAATGCACCCGGGAATCTGGA GGGGCC-3’, rev: 5’-TTATCCACTTCCAATGTTATTAtCCTGGAAAGAGCTTCCGGAATTTGCC-3’). 6xHis-MPB-tagged 43GTPBP6 was expressed in *E.coli* BL21 (DE3) RIL cells (Merck Millipore) grown in LB media. Cells were grown to an optical density at 600 nm of 0.5 at 37 °C and protein expression was subsequently induced with 0.15 mM isopropyl β-d-1-thiogalactopyranoside at 16 °C for 18 h. Cells were collected by centrifugation, resuspended in lysis buffer (300 mM NaCl, 50 mM Na-HEPES pH 7.4, 10% (v/v) glycerol, 30 mM imidazole pH 8.0, 2 mM DTT, 0.284 μg/ml leupeptin, 1.37 μg/ml pepstatin, 0.17 mg/ml PMSF and 0.33 mg/ml benzamidine) and immediately used for protein purification performed at 4 °C. The cells were lysed by sonication and the lysate was cleared by centrifugation (87.200 xg, 4°C, 30 min). The supernatant was applied to a HisTrap HP 5 ml column (GE Healthcare), preequilibrated in lysis buffer. The column was washed with 9.5 CV high-salt buffer (1,000 mM NaCl, 50 mM Na-HEPES pH 7.4, 10% (v/v) glycerol, 30 mM imidazole pH 8.0, 2 mM DTT, 0.284 μg/ml leupeptin, 1.37 μg/ml pepstatin, 0.17 mg/ml PMSF and 0.33 mg/ml benzamidine), and 9.5 CV low-salt buffer (150 mM NaCl, 50 mM Na-HEPES pH 7.4, 10% (v/v) glycerol, 30 mM imidazole pH 8.0 and 2 mM DTT).

The sample was then eluted using nickel elution buffer (150 mM NaCl, 50 mM Na-HEPES pH 7.4, 10% (v/v) glycerol, 500 mM imidazole pH 8.0 and 2 mM DTT). The eluted protein was dialysed overnight in dialysis buffer (150 mM NaCl, 50 mM Na-HEPES pH 7.4, 15 mM imidazole pH 8.0, 10% (v/v) glycerol and 2 mM DTT) in the presence of 4 mg His-tagged TEV protease at 4 °C. The dialysed sample was applied to a HiTrap Heparin HP 5 ml column (GE Healthcare), preequilibrated in Buffer A (20 mM Na-HEPES pH 7.4, 10% (v/v) glycerol, 2 mM DTT) with 7.5 % Buffer B (2 M NaCl, 20 mM Na-HEPES pH 7.4, 10% (v/v) glycerol, 2 mM DTT). The protein was eluted with a linear salt gradient from 7.5 - 50 % Buffer B and peak fractions containing GTPBP6 were collected and re-applied to a His Trap HP 5 ml column in the presence of 40 mM imidazole pH 8.0 to remove cleaved His-tagged MBP and TEV protease. The flow-through containing GTPBP6 was collected and applied to a Superdex 75 10/300 GL column (GE Healthcare) equilibrated in size exclusion buffer (70 mM NH_4_Cl, 30 mM KCl, 7 mM MgCl_2_, 20 mM TRIS-HCl pH 7.4, 10% (v/v) glycerol, 5 mM DTT). Peak fractions were assessed by SDS-PAGE and Coomassie staining and pooled. The protein was concentrated to ~65 mM using a MWCO 30,000 Amicon Ultra Centrifugal Filter (Merck), flash-frozen and stored at −80 °C until use.

### LC-MS/MS analysis

Proteins were separated by polyacrylamide gel electrophoresis on a 4-12% gradient gel (NuPAGE, Invitrogen). After Coomassie staining, lanes were cut into 12 slices and proteins were reduced by dithiothreitol, alkylated by iodoacetamide, and digested with trypsin in-gel. Extracted peptides were vacuum-dried and subsequently resuspended in 2% acetonitrile (ACN, v/v)/0.05% trifluoroacetic acid (TFA, v/v). Peptides were measured on a QExactive HF Mass Spectrometer coupled to a Dionex UltiMate 3000 UHPLC system (both Thermo Fisher Scientific) equipped with an in house-packed C18 column (ReproSil-Pur 120 C18-AQ, 1.9 μm pore size, 75 μm inner diameter, 30 cm length, Dr. Maisch GmbH). Peptides were separated applying the following gradient: mobile phase A consisted of 0.1% formic acid (FA, v/v), mobile phase B of 80% ACN/0.08% FA (v/v). The gradient started at 5% B, increasing to 10% B within 3 min, followed by a continuous increase to 46% B within 45 min, and then keeping B constant at 90% for 8 min. After each gradient the column was again equilibrated to 5% B for 2 min. The flow rate was set to 300 nL/min. MS1 full scans were acquired with a resolution of 60,000, an injection time (IT) of 50 ms and an automatic gain control (AGC) target of 1×10^6^. Dynamic exclusion (DE) was set to 30 s. MS2 spectra were acquired of the 30 most abundant precursor ions; the resolution was set to 15,000; the IT was set to 60 ms and the AGC target to 1×10^5^. Fragmentation was enforced by higher-energy collisional dissociation (HCD) at 28% NCE. Acquired raw data were analyzed by MaxQuant^43^ (v. 1.6.0.1) applying default settings and enabled ‘match between runs’ option. Proteins were quantified based on their iBAQ value.

The mass spectrometry proteomics data have been deposited to the ProteomeXchange Consortium via the PRIDE ^44^ partner repository with the dataset identifier PXD023502.

### Bisulfite sequencing to monitor 12S-m^5^C1488 and 12S-m^4^C1486

To ensure that the accumulation of NSUN4 on the mtLSU in GTPBP6-deficient cells does not affect its second function as a methyltransferase modifying the 12S rRNA at position 1488, we analyzed the modification status of 12S-m^5^C1488 and, as a control, 12S-m^4^C1486, by subjecting DNase-treated total RNA from HEK293T wild type and *Gtpbp6*^−/−^ cells to bisulfite sequencing ^45,46^. Bisulfite treatment was performed using the EpiTect Bisulfite kit (Qiagen) according to the manufacturer’s instructions. Deamination was performed by three cycles of incubation at 70°C for 5 min and at 60°C for 60 min. Samples were purified using mini Quick spin RNA columns (Roche) and the desulphonated in Tris pH 9.0 for 30 min at 37°C. RNA was extracted using phenol:chloroform, precipitated and reverse transcribed from the 12S-m^5^C841_RT primer (5’-TTTAATTAAATATCCTTTAAAATATAC-3’) using Superscript III reverse transcriptase (Thermo) according to the manufacturer’s instructions. A 70 nt fragment of the 12S rRNA was amplified by PCR (5’-TTTAATTAAATATCCTTTAAAATATAC-3’; 5’-AATAGGGTTTTGAAGTGTGTATATA-3’) and cloned using a TOPO-TA kit (Thermo). Clones were sequenced at Eurofins Genomics using the T7 primer and only sequences in which all cytosines (disregarding C1486 and C1488) were converted to uracil/thymine were used for the presented analysis (**Extended Data Fig. 7c**).

## EXTENDED DATA FIGURES

**Extended Data Fig. 1.**
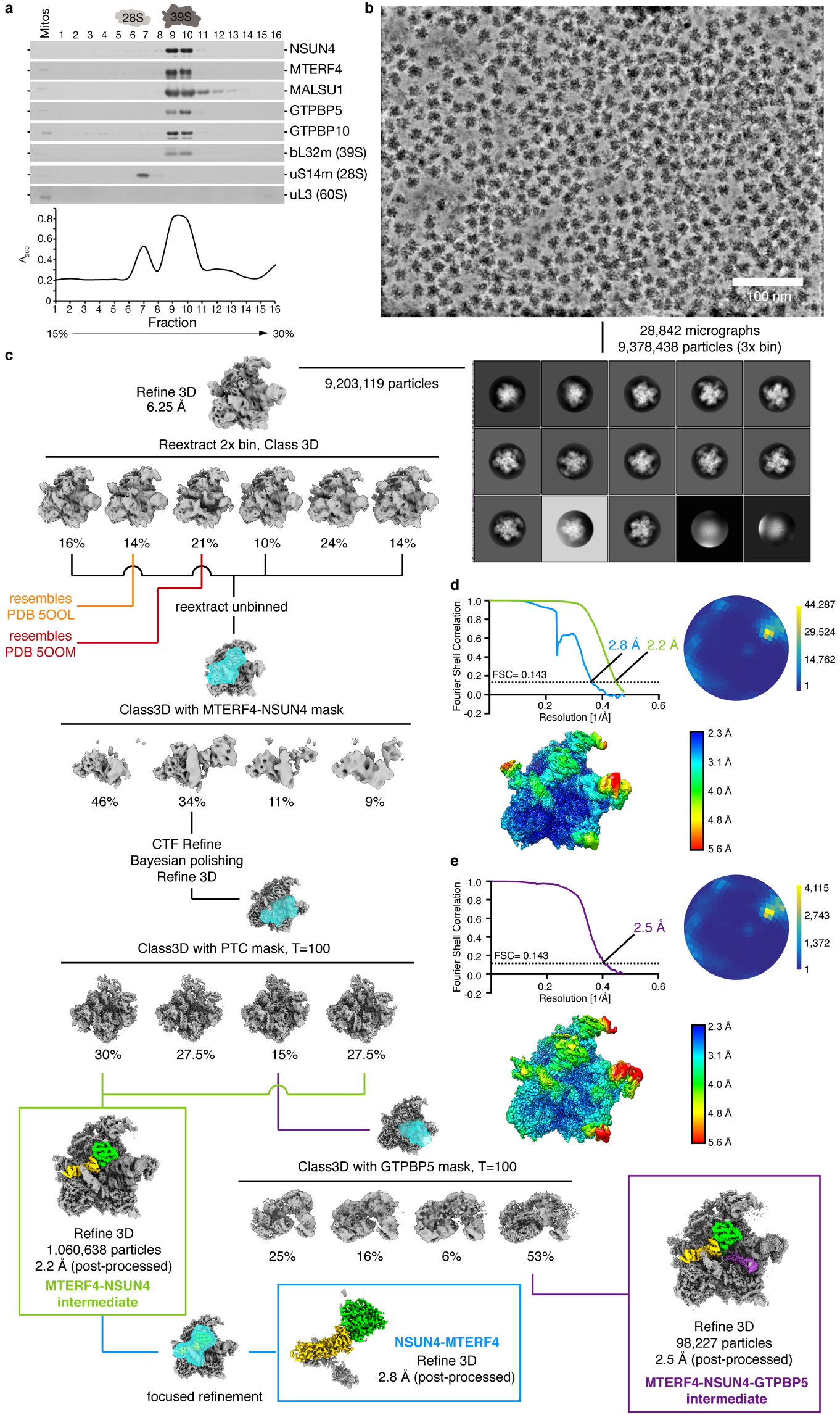
Purification and Cryo EM data processing first dataset. **(a)** Gradient fractions (1-16) upon ribosome isolation were analyzed by western blotting using indicated antibodies. uL3 (60S LSU) was used as a control to assess the level of contamination with cytosolic ribosomes. Absorbance was measured for each fraction at 260 nm. Fractions 8 and 9 were used for further analyses. **(b)** Example denoised micrograph calculated from two independently measured half sets of 40 frames each. Scale bar, 100 nm. **(c)** 2D class averages and Cryo-EM processing tree. **(d)** Fourier shell correlation (FSC) plot, Angular distribution plot and local resolution distribution for the MTERF4-NSUN4 mtLSU intermediate. Scale for the angular distribution plot shows the number of particles assigned to a particular angular bin. Blue, a low number of particles; yellow, a high number of particles. **(e)** Fourier shell correlation (FSC) plot, Angular distribution plot and local resolution distribution for the MTERF4-NSUN4-GTPBP5 mtLSU intermediate. Scale for the angular distribution plot shows the number of particles assigned to a particular angular bin. Blue, a low number of particles; yellow, a high number of particles.

**Extended Data Fig. 2.**
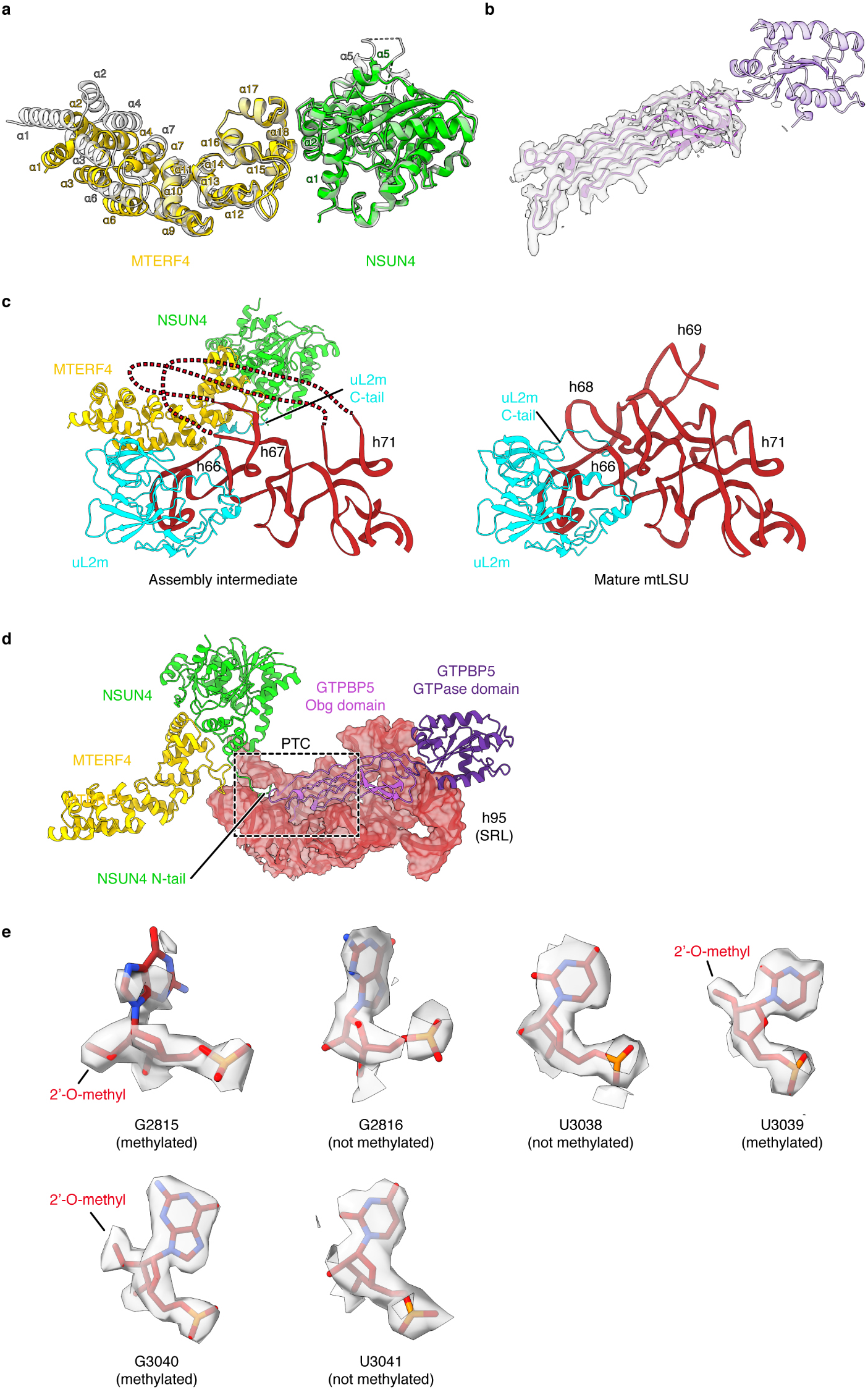
Structural details of MTERF4, NSUN4 and ribosomal RNA. **(a)** Comparison between the MTERF4-NSUN4 complex bound to the mtLSU and the previous crystal structure (PDB 4FP9) ^15^. The two structures are shown superimposed as cartoon. The mtLSU-bound structure is colored as in Figure 1 and the free crystal structure is colored in grey and shown transparently. Secondary structure elements in MTERF4 and that adopt different conformations in NSUN4 are indicated. The helical repeats of MTERF4 form a widened curve when bound to the ribosome. **(b)** Density fit of GTPBP5. The model of GTPBP5 is shown as cartoon with coloring as in Figure 1. The cryo-EM density (before postprocessing) is shown as transparent grey surface. **(c)** Close-up view of the MTERF4-NSUN4 complex bound to the mtLSU as shown in Figure 1 (left). Close-up view of the 16S rRNA region 2469-2659 in the mature mtLSU (PDB 3J7Y)^3^ (right). The fold observed in the mature mtLSU would clash with MTERF4-NSUN4. **(d)** Interaction of GTPBP5 with the mtLSU. Regions of the 16S rRNA interacting with MTERF4-NSUN4 are shown as cartoon and as transparent surface. MTERF4, NSUN4 and GTPBP5 are shown as cartoons. **(e)** G2815, U3039 and G3040 are methylated. G2815, U3039 and G3040 are shown as sticks with the postprocessed cryo-EM density of the MTERF4-NSUN4-GTPBP5 mtLSU intermediate (dataset 1) shown as transparent surface. Neighboring non-methylated bases are shown for comparison.

**Extended Data Fig. 3.**
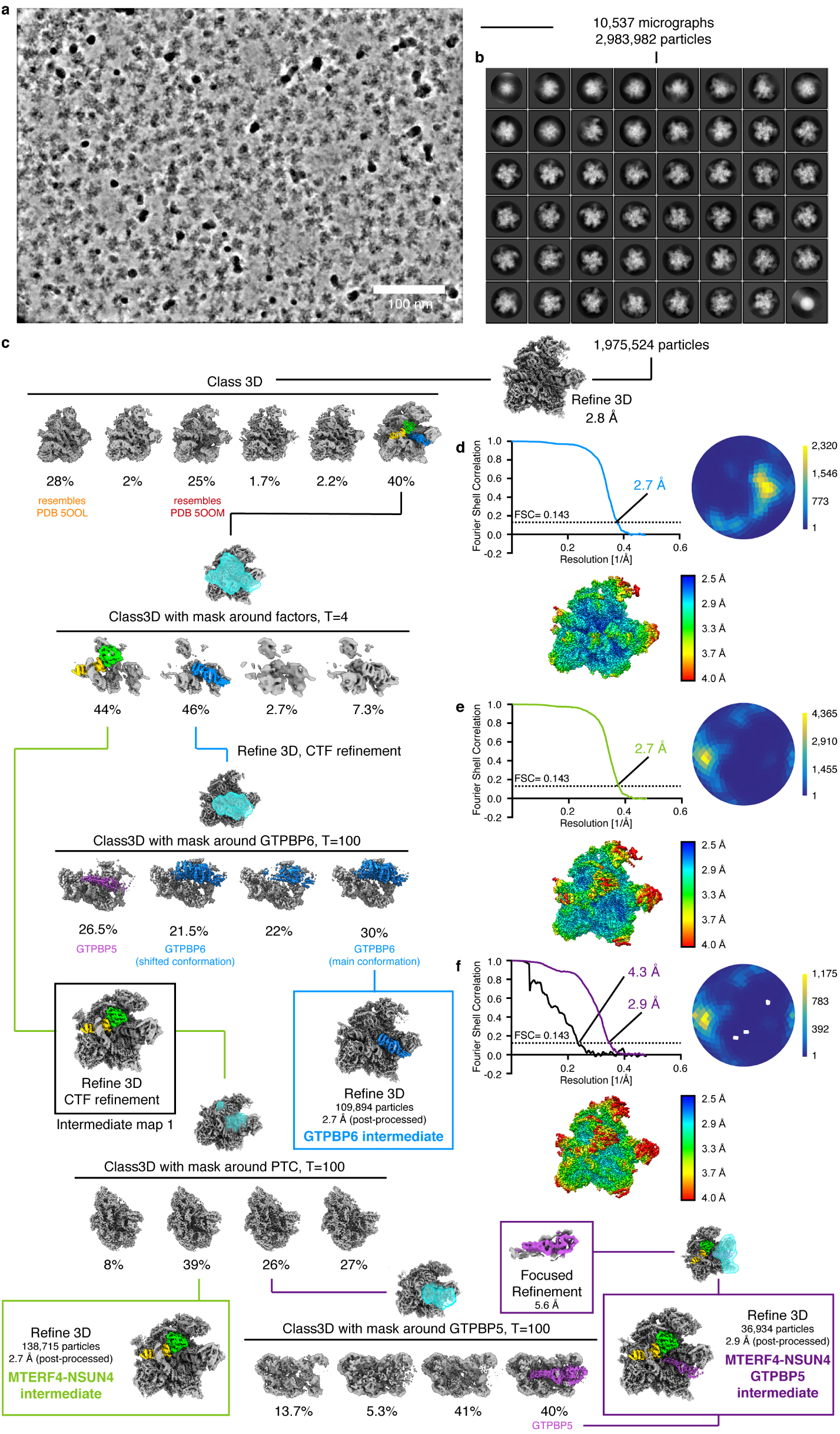
Cryo-EM data processing second dataset. **(a)** Example denoised micrograph calculated from two independently measured half sets of 40 frames each. Scale bar, 100 nm. **(b)** 2D class averages. **(c)** Cryo-EM processing tree. **(d)** Fourier shell correlation (FSC) plot, Angular distribution plot and local resolution distribution for the GTPBP6-bound mtLSU intermediate. Scale for the angular distribution plot shows the number of particles assigned to a particular angular bin. Blue, a low number of particles; yellow, a high number of particles. **(e)** Fourier shell correlation (FSC) plot, Angular distribution plot and local resolution distribution for the MTERF4-NSUN4 mtLSU intermediate. Scale for the angular distribution plot shows the number of particles assigned to a particular angular bin. Blue, a low number of particles; yellow, a high number of particles. **(f)** Fourier shell correlation (FSC) plot, Angular distribution plot and local resolution distribution for the MTERF4-NSUN4-GTPBP5 mtLSU intermediate. Scale for the angular distribution plot shows the number of particles assigned to a particular angular bin. Blue, a low number of particles; yellow, a high number of particles.

**Extended Data Fig. 4.**
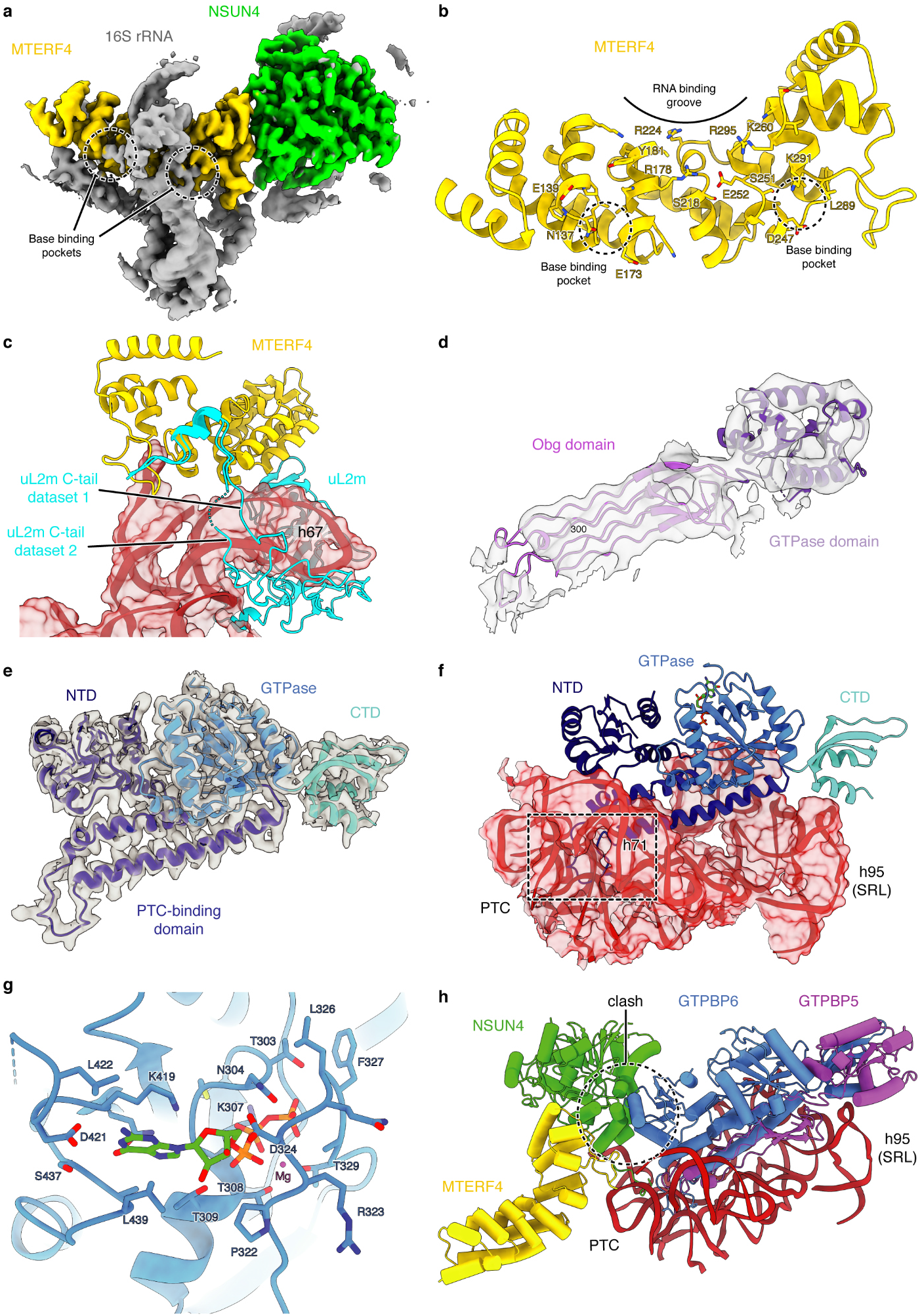
Structural details of MTERF4, NSUN4, GTPBP5 and GTPBP6 in the second dataset. **(a)** Improved density for the 16 rRNA region wrapping over MTERF4. A region of the cryo-EM reconstruction (Intermediate map 1, **Extended Data Fig. 3**) of MTERF4-NSUN4 containing particles from dataset 2 is shown as surface and colored as follows: NSUN4: lime green, MTERF4: yellow, 16S rRNA: grey. The region of the rRNA wrapping above MTERF4 adopts a helical fold and may form base-mediated interactions with MTERF4 (indicated). **(b)** Close-up view of the RNA-binding groove of MTERF4. MTERF4 is shown as cartoon in yellow and residues close to the RNA density observed in (a) are shown as sticks. The potential base-binding pockets are indicated. **(c)** The C-tail of uL2m occupies different paths in the MTERF4-NSUN4 mtLSU intermediate from dataset 1 and dataset 2. MTERF4 and uL2m are shown as cartoons and the 16S rRNA elements interacting with them are shown as cartoon and transparent surface. Coloring as in Figure 1. **(d)** Improved density for the GTPBP5 GTPase domain in dataset 2. GTPBP5 is shown as cartoon and colored as in Figure 1. The cryo-EM density obtained from focused refinement of the GTPBP5-containing particle set in dataset 2 is shown as transparent grey surface. **(e)** Density fit of GTPBP6. GTPBP6 is shown as cartoon and colored as in Figure 3. The cryo-EM density (not post-processed) of the GTPBP6-bound mtLSU intermediate structure is shown as transparent grey surface. **(f)** Interaction of GTPBP6 with the mtLSU. Regions of the 16S rRNA interacting with GTPBP6 are shown as cartoon and as transparent surface. GTPBP6 is shown as cartoon. **(g)** GTP binding pocket of GTPBP6. The GTP binding site of GTPBP6 is shown as cartoon and colored as in Figure 3. Residues within 4 Å of GTP are shown as sticks. GTP is shown in green as sticks. **(h)** GTPBP6 occupies the same binding site as GTPBP5 on the mtLSU and would clash with NSUN4. The structure of the MTERF4-NSUN4-GTPBP5 intermediate and the GTPBP6 intermediate were superimposed with their 16S rRNA. The PTC and factors are shown as cartoons and colored as in Figures 1 and 3.

**Extended Data Fig. 5.**
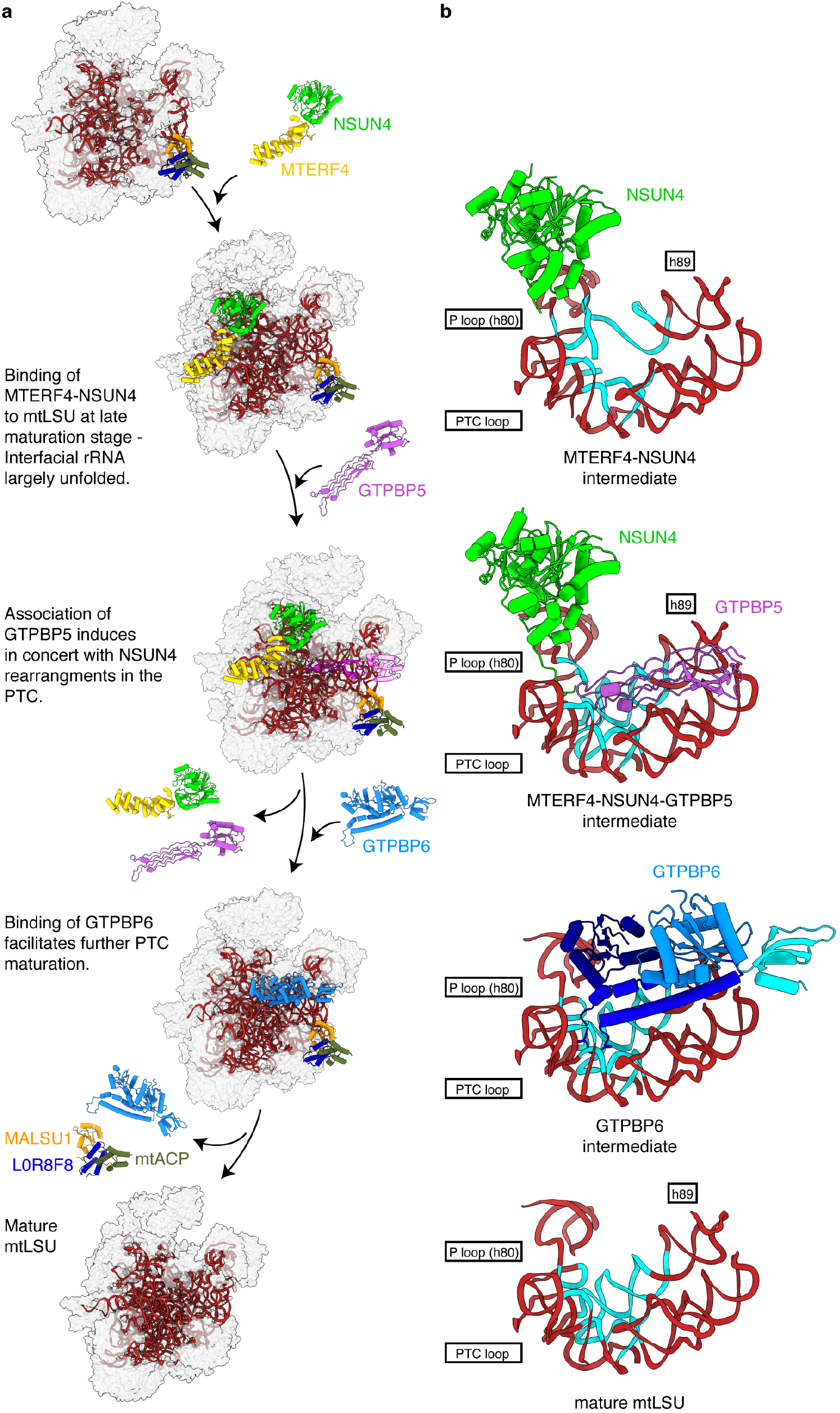
Maturation of human mtLSU. **(a)** Intermediate states of mtLSU are depicted as in Figure 1a and 3a. Structures of immature and mature mtLSU were used from previous studies (PDB 5OOM, 3J7Y) ^3,5^. **(b)** Model for GTPase-mediated PTC maturation. PTC intermediate states are depicted as in Figure 2 and Extended Data Figure 5. The mature mtLSU was modeled based on a previous structure (PDB 3J7Y)^3^.

**Extended Data Fig. 6.**
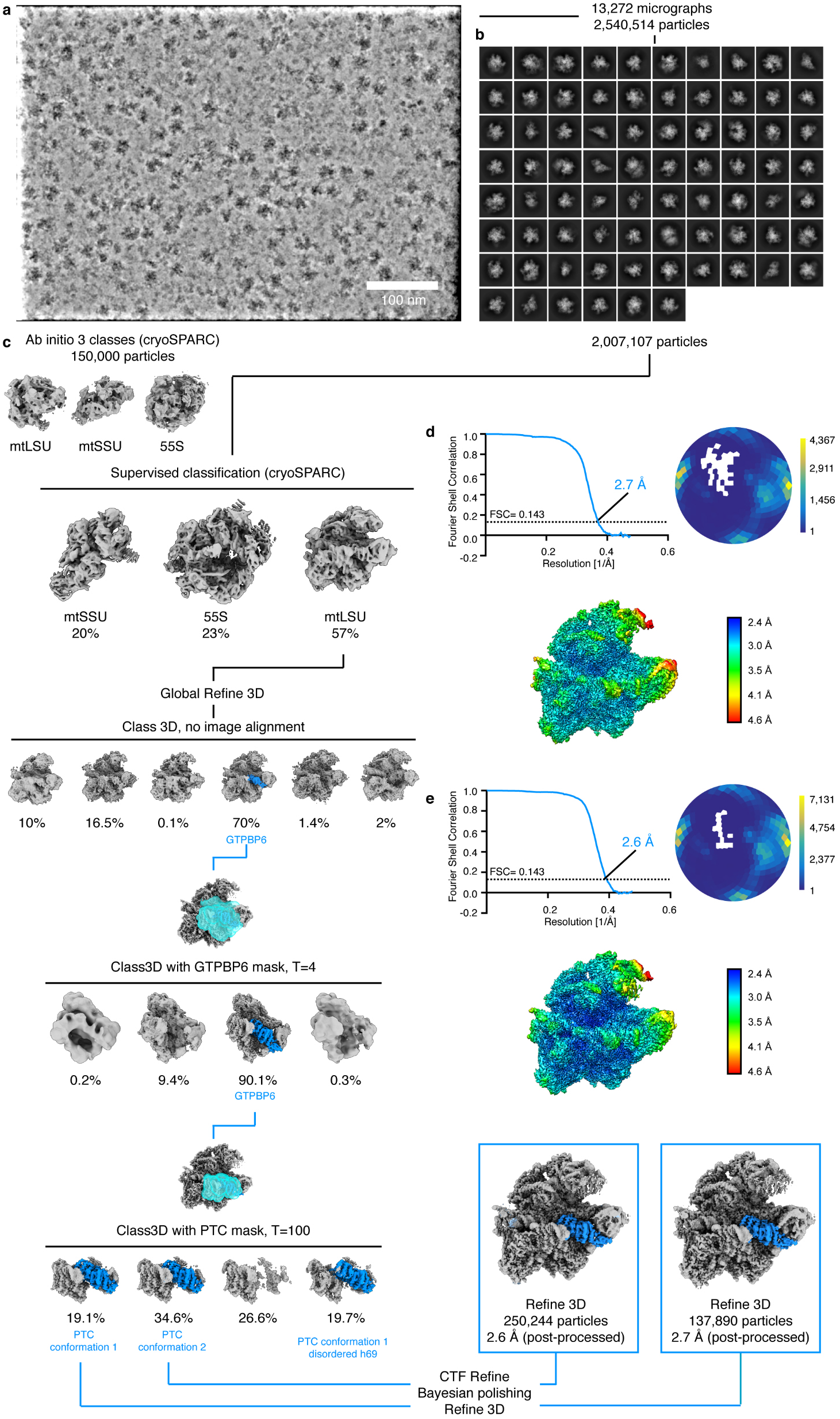
Cryo-EM data processing third dataset. **(a)** Example denoised micrograph calculated from two independently measured half sets of 40 frames each. Scale bar, 100 nm. **(b)** 2D class averages. **(c)** Cryo-EM processing tree. **(d)** Fourier shell correlation (FSC) plot, Angular distribution plot and local resolution distribution for the GTPBP6-bound split mtLSU with PTC conformation 1. Scale for the angular distribution plot shows the number of particles assigned to a particular angular bin. Blue, a low number of particles; yellow, a high number of particles. **(e)** Fourier shell correlation (FSC) plot, Angular distribution plot and local resolution distribution for the GTPBP6-bound split mtLSU with PTC conformation 2. Scale for the angular distribution plot shows the number of particles assigned to a particular angular bin. Blue, a low number of particles; yellow, a high number of particles.

**Extended Data Fig. 7.**
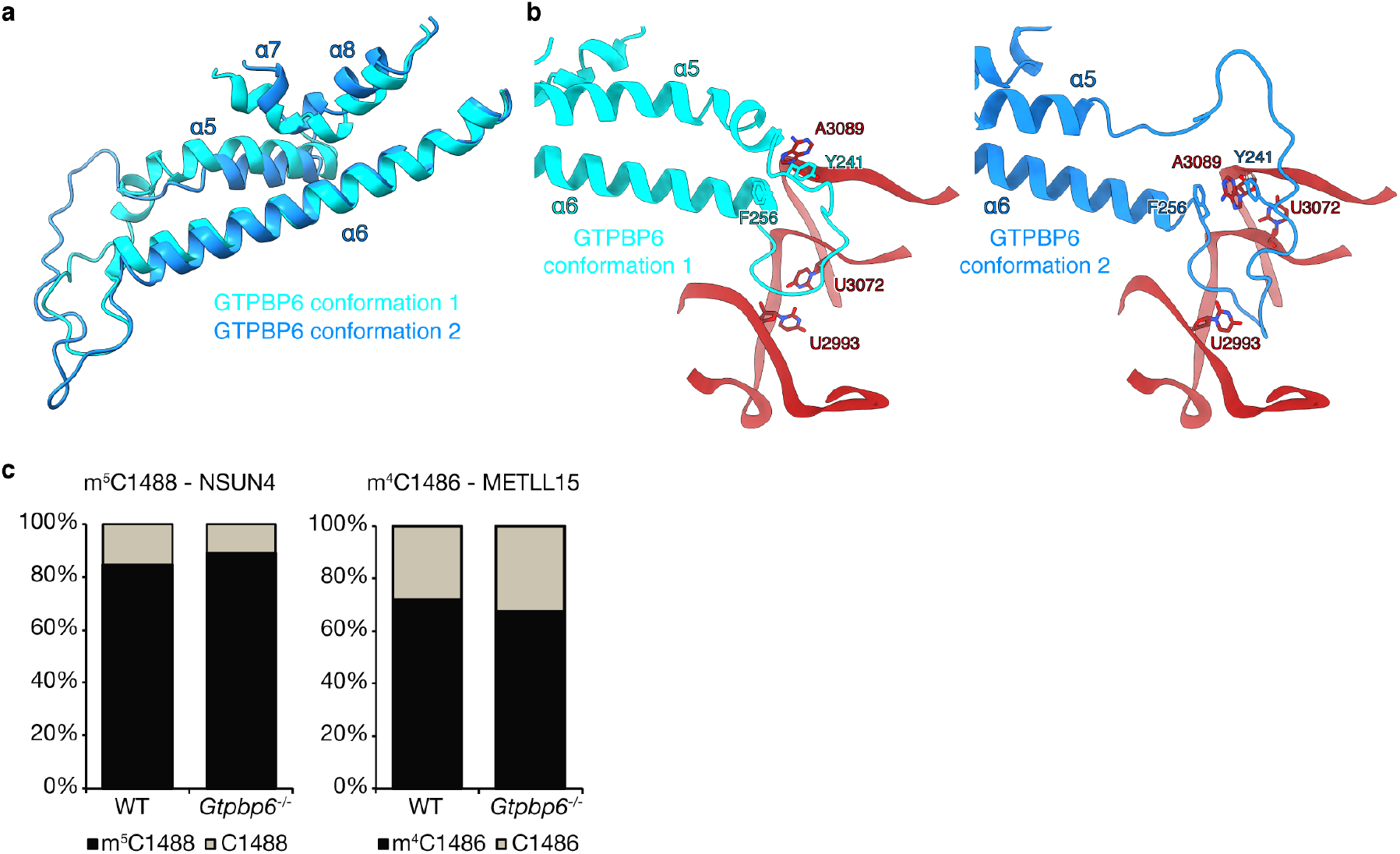
GTPBP6 conformations. **(a)** The PTC-binding loop of GTPBP6 can adopt two conformations. Superimposition of GTPBP6 in the PTC conformation 1 and PTC conformation 2 structures observed after ribosome splitting. GTPBP6 is shown as cartoon. GTPBP6 conformation 1, which is also observed during mtLSU biogenesis, is shown in cyan and GTPBP6 conformation 2 in marine. **(b)** Close-up of PTC interactions in GTPBP6 PTC conformation 1 and 2. Depiction as in a, with the region of the 16S rRNA differing between the two states shown as cartoon. Bases that adopt different conformations as well as interacting GTPBP6 residues are shown as sticks. **(c)** Methylation of 12S rRNA is not affected in GTPBP6-deficient cells. Total RNA from wild type (WT) or *Gtpbp6*^−/−^ was treated with bisulfite, reverse transcribed and a region of the 16S rRNA containing m^5^C1488 and m^4^C1486 was amplified, cloned and sequenced. The relative proportions of unconverted cytosine reflecting m^5^C/m^4^C (black) and thymine reflecting converted, unmodified cytosine (grey) are shown. Data from >45 clones is presented.

## EXTENDED DATA TABLES

**Extended Data Table 1.** Cryo-EM data collection and refinement for dataset 1.

|  | **MTERF4-NSUN4**<br>**intermediate** | **MTERF4-NSUN4-**<br>**GTPBP5-intermediate** |
| --- | --- | --- |
| Data collection and processing |  |  |
| Magnification | 81,000 | 81,000 |
| Voltage (kV) | 300 | 300 |
| Electron exposure (e-/Å<sup>2</sup>) | 36 | 36 |
| Defocus range (μm) | 0.3 – 2.8 | 0.3 – 2.8 |
| Pixel size (Å) | 1.05 | 1.05 |
| Symmetry imposed | C1 | C1 |
| Initial particle images (no.) | 9,378,438 | 9,378,438 |
| Final particle images (no.) | 1,060,638 | 98,227 |
| Map resolution (Å) | 2.2 | 2.5 |
| FSC threshold | 0.143 | 0.143 |
| Map resolution range (Å) | 5.6 – 2.2 | 5.6 – 2.2 |
| Refinement |  |  |
| Model resolution (Å) | 2.3 | 2.7 |
| FSC threshold | 0.5 | 0.5 |
| Map sharpening *B* factor (Å<sup>2</sup>) | -50 | -39.5 |
| Model composition |  |  |
| Non-hydrogen atoms | 101,807 | 101,495 |
| Protein/Nucleotide residues | 8,685 | 8,704 |
| Ligands | ZN:3, UNK:28, MG:52 | Zn:3, UNK:28, MG:52 |
| *B* factors (Å<sup>2</sup>) |  |  |
| Protein/Nucleotide | 106.96 / 97.55 | 93.43 / 73.19 |
| Ligand | 141.25 | 153.64 |
| R.m.s. deviations |  |  |
| Bond lengths (Å) | 0.009 | 0.010 |
| Bond angles (°) | 0.859 | 0.898 |
| Validation |  |  |
| MolProbity score | 2.16 | 1.94 |
| Clashscore | 14.35 | 14.30 |
| Poor rotamers (%) | 2.19 | 0.05 |
| Ramachandran plot |  |  |
| Favored (%) | 96.47 | 95.98 |
| Allowed (%) | 3.51 | 3.95 |
| Disallowed (%) | 0.02 | 0.07 |

**Extended Data Table 2.** Cryo-EM data collection and refinement for dataset 2.

|  | **GTPBP6**<br>**intermediate** | **MTERF4-NSUN4**<br>**intermediate** | **MTERF4-**<br>**NSUN4-**<br>**GTPBP5-**<br>**intermediate** |
| --- | --- | --- | --- |
| Data collection and processing |  |  |  |
| Magnification | 81,000 | 81,000 | 81,000 |
| Voltage (kV) | 300 | 300 | 300 |
| Electron exposure (e-/Å<sup>2</sup>) | 37 | 37 | 37 |
| Defocus range (μm) | 0.3 – 2.1 | 0.3 – 2.1 | 0.3 – 2.1 |
| Pixel size (Å) | 1.05 | 1.05 | 1.05 |
| Symmetry imposed | C1 | C1 | C1 |
| Initial particle images (no.) | 2,983,982 | 2,983,982 | 2,983,982 |
| Final particle images (no.) | 109,894 | 138,715 | 36,934 |
| Map resolution (Å) | 2.7 | 2.7 | 2.9 |
| FSC threshold | 0.143 | 0.143 | 0.143 |
| Map resolution range (Å) | 4.0 – 2.5 | 4.0 – 2.5 | 4.0 – 2.5 |
| Refinement |  |  |  |
| Model resolution (Å) | 2.8 |  |  |
| FSC threshold | 0.5 |  |  |
| Map sharpening *B* factor (Å<sup>2</sup>) | -52 |  |  |
| Model composition |  |  |  |
| Non-hydrogen atoms | 101,494 |  |  |
| Protein/Nucleotide residues | 8,616 |  |  |
| Ligands | GTP:1, ZN:3,<br>UNK:28,<br>MG:52 |  |  |
| *B* factors (Å<sup>2</sup>) |  |  |  |
| Protein/Nucleotide | 80.36 / 59.19 |  |  |
| Ligand | 110.29 |  |  |
| R.m.s. deviations |  |  |  |
| Bond lengths (Å) | 0.007 |  |  |
| Bond angles (°) | 0.730 |  |  |
| Validation |  |  |  |
| MolProbity score | 1.82 |  |  |
| Clashscore | 11.67 |  |  |
| Poor rotamers (%) | 0.01 |  |  |
| Ramachandran plot |  |  |  |
| Favored (%) | 96.45 |  |  |
| Allowed (%) | 3.51 |  |  |
| Disallowed (%) | 0.04 |  |  |

**Extended Data Table 3.**
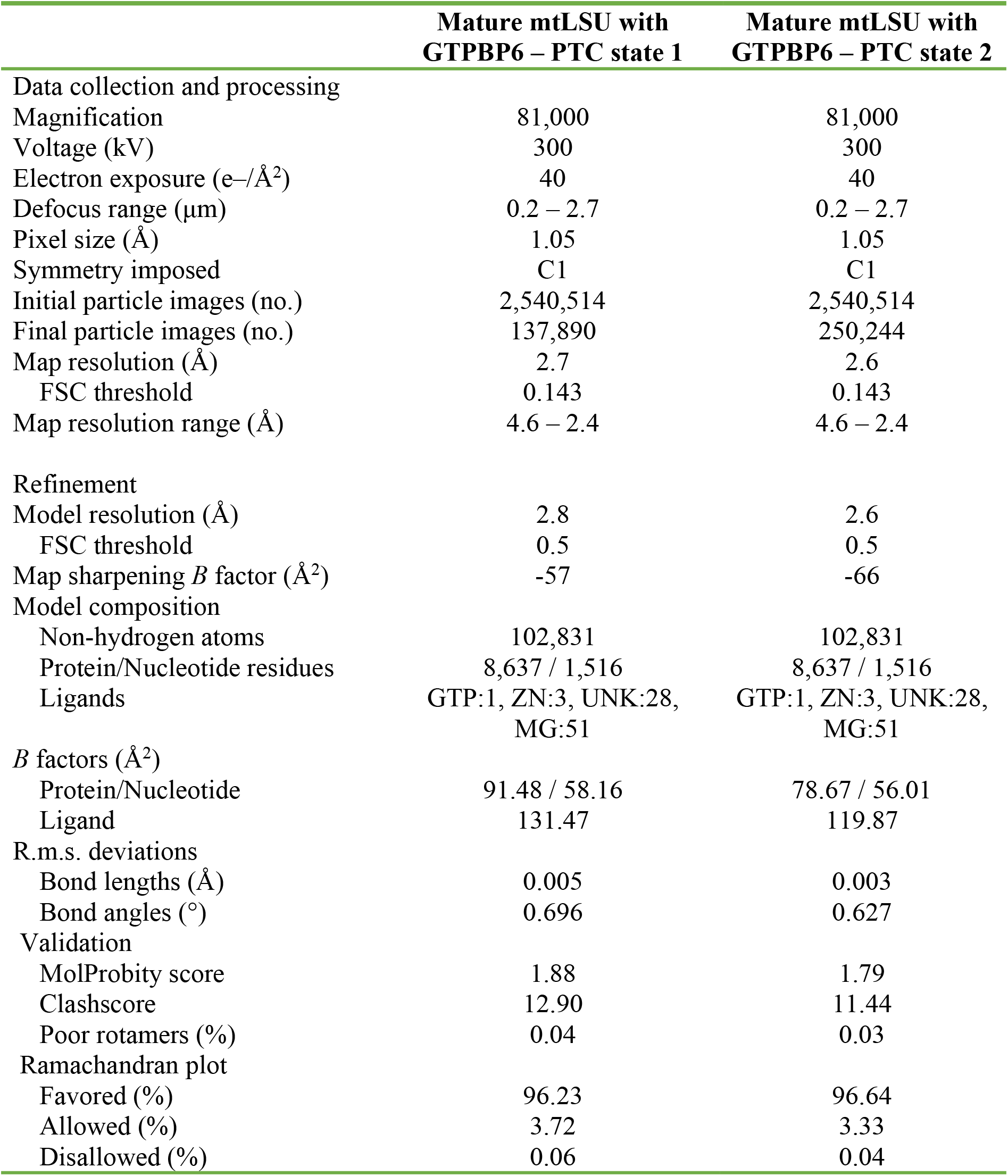
Cryo-EM data collection and refinement for dataset 3.

## SUPPLEMENTARY MATERIALS

**Supplementary Table 1 | LC-MS/MS analysis of mtLSU and mtSSU purified from GTPBP6-deficient cells.**

**Supplementary Video 1 | Model of peptidyl transferase center maturation during human mitochondrial ribosome biogenesis.**

